# Targeting USP9x/SOX2 axis contributes to the anti-osteosarcoma effect of neogambogic acid

**DOI:** 10.1101/714741

**Authors:** Xiangyun Chen, Xingming Zhang, Haiyan Cai, Wupeng Yang, Hu Lei, Hanzhang Xu, Weiwei Wang, Qi Zhu, Jingwu Kang, Tong Yin, Wenli Gu, Ying-Li Wu

## Abstract

SOX2 has been viewed as a critical oncoprotein in osteosarcoma. Emerging evidence show that inducing the degradation of transcription factors such as SOX2 is a promising strategy to make them druggable. Here, we show that neogambogic acid (NGA), an active ingredients in *garcinia*, significantly inhibited the proliferation of osteosarcoma cells with ubiquitin proteasome-mediated degradation of SOX2 *in vitro* and *in vivo*. We further identified USP9x as a bona fide deubiquitinase for SOX2 and NGA directly interacts with USP9x in cells. Moreover, knockdown of USP9x inhibited the proliferation and colony formation of osteosarcoma cells, which could be rescued by overexpression of SOX2. Consistent with this, knockdown of USP9x inhibited the proliferation of osteosarcoma cells in a xenograft mouse model. Collectively, we identify USP9x as the first deubiquitinating enzyme for controlling the stability of SOX2 and USP9x is a direct target for NGA. We propose that targeting the USP9x/SOX2 axis represents a novel strategy for the therapeutic of osteosarcoma and other SOX2 related cancers.

## Introduction

SOX2 (SRY-Related HMG-Box Gene 2), one of the SoxB1 subfamily transcription factors, is essential for embryonic stem cell maintenance, pluripotency and self-renewal (Feng & Wen, 2015). SOX2 can reprogram differentiated cells into pluripotent cells cooperating with Oct4, KLF4 and c-Myc. Overexpression of SOX2 is involved in the development of various cancers such as ovarian cancer, and hepatocellular carcinoma (Liu et al, 2013). Recently, SOX2 is reported to play a critical role in osteosarcoma, the most common primary malignant bone cancer (Maurizi et al, 2018b; Wang et al, 2017). SOX2 is highly expressed in human osteosarcoma cell lines as well as in the tumor samples, and is essential for osteosarcoma cell self-renewal (Basu-Roy et al, 2012). SOX2 is able to antagonize the Hippo pathway to maintain an undifferentiated state of osteosarcoma (Basu-Roy et al, 2015; Maurizi et al, 2018a). Therefore, SOX2 is an attractive therapeutic target for osteosarcoma. However, unlike protein kinases, transcription factors like SOX2 are commonly considered as undruggable. Encouragingly, inducing degradation of transcription factors such as PML-RARα or IKZF1 has been successful used in the treatment of leukemia and myeloma. This indicates that inducing degradation of SOX2 maybe an alternative strategy to target SOX2 (Kronke et al, 2014). Thus, it is important to dissect the mechanisms regulating the degradation of SOX2 in osteosarcoma (Huser et al, 2018). The ubiquitin-proteasome system has been involved in controlling the stability of SOX2. Several E3 ubiquitin ligases for SOX2 has been reported (Cicconi et al, 2018; Wang et al, 2016). However, the deubiquitinating enzymes for controlling SOX2 stabiligy remains unknown.

USP9x, the homolog of *Drosophila fat facets* (Wood et al, 1997), belongs to the ubiquitin specific proteases subfamily group of deubiquitinases. Interestingly, USP9x can play a dual role in cancers, promoting or suppressing tumor development, depending on the cancer type and disease model studied. For example, by stabilizing MCL1, USP9x can promote cancer cell survival or confer radioresistance in B-cell acute lymphoblastic leukemia (Zhou et al, 2015), non-small-cell lung cancer (Yan et al, 2014) breast (Li et al, 2017) and melanoma (Potu et al, 2017a). However, in pancreatic ductal adenocarcinoma, loss of USP9x enhances transformation and protects pancreatic cancer cells from apoptosis (Perez-Mancera et al, 2012). In colorectal cancer, USP9x can suppresses proliferation of cancer cells through stabilizing FBW7 (Khan et al, 2018b). In osteosarcoma, the role of USP9x remains unknown.

Neogambogic acid (NGA) is one of the active ingredients in *garcinia* (Liu et al, 2015a; Ranjbarnejad et al, 2017; Sethi et al, 2014; Zhou et al, 2017). Preliminary studies show that the neogambogic acid can selectively inhibit the growth of various cancer cells, and has a broader antitumor activity and lower toxicity than gambogic acid. NGA can induce apoptosis in colon cancer, breast cancer, melanoma, hepatoma, myeloma and leukemia. The anti-cancer effect has been attribute to the inhibition of the MAPK signaling pathway, or the activity of telomerase or inducing endoplasmic reticulum stress (Sun et al, 2018; Wang et al, 2011). However, the precision target of NGA is not known.

In this study, we demonstrate that NGA can inhibit the proliferation of osteosarcoma cells *in vitro* and *in vivo*, in which the reduction of SOX2 plays an important role. Moreover, we identify USP9x as the first deubiquitinating enzyme for controlling the stability of SOX2 and USP9x is a direct target for NGA. We propose that targeting the USP9x/SOX2 axis represents a novel strategy for the therapeutic of osteosarcoma.

## Materials and Methods

### Cell culture

The human osteosarcoma cell lines U2OS and 143B were maintained in Dulbecco’s minimum essential medium (DMEM, Hyclone, USA) that was supplemented with 10% fetal bovine serum (FBS, Gibco BRL, USA), 100 units/ml penicillin, and 100 lg/ml streptomycin, at 37°C in a humidified atmosphere containing 5% CO_2_.

### Western blot analysis

Cells were collected and lysed with lysis buffer (50 mM TrisHCl, pH 6.8, 100 mM DTT, 2% SDS, 10% glycerol), then protein through denaturation with high temperature and centrifuged at 20,000g for 10 min, and protein in the supernatants were quantified. Protein extracts were equally loaded on 8%–12% SDS polyacrylamide gel, electrophoresed, and transferred to a nitrocellulose membrane (Bio-Rad, CA). The blots were stained with 0.2% Ponceau red S to ensure equal protein loading. After blocked with 5% nonfat milk in PBS for 1 hour, the membranes were probed with antibodies overnight. According to the manufacturer’s instructions, the signals were detected with a chemiluminescence phototope-HRP kit (Cell Signaling, USA). As necessary, blots were stripped and reprobed with anti-β-actin (Calbiochem, Germany) as internal control. All experiments were repeated three times.

### Cell proliferation assay

Inhibition of cell proliferation induced by NGA was measured by Cell Counting Kit-8 assay kit (Dojindo, Japan). Cells (1 × 10^4^/well) were seeded into a 96-well plate and grown overnight, then treated with different concentrations of NGA (SELLECK, USA). After incubation for 24 h or 48h, 10 μl of CCK8 reagent was added to each well. After incubation for another 2 h, the absorbance at 450 nm was measured using Synergy H4 Hybrid Microplate Reader (Synergy H4, USA). The cell proliferation inhibition ratio was calculated by the following formula: cell proliferation inhibition ratio (%) = (O.D. control – O.D. treated)/O.D. control × 100%.

### Cell cycle and apoptosis assays

Propidium iodide (PI, Sigma) staining was used to analyze cell cycle distribution. After exposure to 1 μM NGA for 24 h or knockdown of USP9x, cells were harvested and fixed with 70% ethanol at −20 °C overnight. Cells were then washed twice with pre-cooled phosphate-buffered saline (PBS), incubated with PBS containing 10 μl RNase A (25 μg/ml) at 37 °C for 30 min, and stained with PI (1 mg/ml in PBS) for 15 min. The DNA content of cells and cell cycle distribution were analyzed by Flow cytometric analysis on BD LSRFortessa cell analyzer (BD Biosciences).

The apoptosis rate of cells was determined using Annexin V-APC and PI (BD Pharmingen) staining. Osteosarcoma cells (5 × 10^5^/ml) in 6-well plates were treated with different concentrations of NGA for 24 h. Cells were collected, washed with PBS twice and 1 × binding buffer once, then resuspended in 400 μl of 1 × binding buffer. Next, 5 μl of Annexin V-APC and 5 μl of PI were added to each sample, and the samples were incubated in the dark for 15 min. Cells were analyzed by fluorescence-activated cell sorting using a BD LSRFortessa cell analyser (BD Biosciences), and the results were analyzed using FlowJo 7.0 software (Tree Star, Ashland, OR, USA).

### Cell clonogenic assay

Cells were seeded into 6-well plates at a density of 2 × 10^3^ cells/well in 2 ml medium containing 10% FBS. Culture medium was changed every 3 days for 2 weeks. The cell clones were stained for 15 min with the solution containing 0.5% crystal violet and 25% methanol, followed by rinsing with tap water three times to remove excess dye. Colonies consisting of more than 50 cells were counted under microscope.

### Cellular thermal shift assay (CETSA)

1 × 10^7^ U2OS cells were collected and washed with ice-cold PBS three times. The cells were resuspended with 1 ml of ice-cold PBS with Roche complete EDTA-free protease inhibitor cocktail (1:100), followed by three snap-freeze cycles consisting of 30 sec to 1 min in liquid nitrogen and then at 25 °C in a thermal cycler or heating block until thawed. Cell lysates were then centrifuged at 20,000 g for 20 min at 4 °C to pellet cellular debris. To determine melting curves, cell lysates were divided into two aliquots; one was treated with NGA and the other with the corresponding concentration of DMSO (control). After 30 min of incubation at room temperature, the lysates were divided into smaller (35 μl) aliquots and heated individually at different temperatures (47.1, 50.3, 54.4, 59.9, 64.2, 67.1 or 69 °C) for 3 min, followed by cooling for 3 min at room temperature. The heated lysates were centrifuged at 20,000 g for 20 min at 4 °C in order to separate the soluble fractions from precipitates. The supernatants were transferred to new microtubes and analyzed by SDS-PAGE followed by western blots. The procedure for establishing isothermal dose response curves was similar to that for melting curves, except that the compound concentration rather than temperature was varied. All cells were heated at 52 °C, which was determined based on analysis of the data obtained during melting curve experiments.

### Molecular docking

The binding modes between USP9X and NGA were analyzed using software AutoDock Vina. The X-ray crystal structure of USP9X was retrieved from protein data bank (PDB code: 5WCH) for docking simulation. All crystallographic water were removed. The compound was docked to a binding pocket, which was centered on the coordinate center of N1829. The binding site was covered by preparing a 60×60×60 size of grid box with grid spacing of 0.375Å of spacing between grid points. The top ranked conformation of the compound was selected for binding mode analysis.

### RNA isolation and quantitative real-time PCR

Total RNA from U2OS and 143B cells was extracted using TRIZOL Reagent (Invitrogen, USA) and cDNA synthesis was performed using the RT Kit (TransGen Biotech, China). Then gene expression of GAPDH, USP9x, SOX2, was detected using Power SYBR Green PCR master mix (Roche, Switzerland). Data were collected and analyzed quantitatively on ABI Prism 7500 sequence detection system (Life Technologies Corporation, USA). GAPDH was used as endogenous control to normalize the differences of total RNA in each sample. The primer sequences used were as follows: for GAPDH (F: 5’-TTGGTATCGTGGAAGGACTC-3’, R: 5’-ACAGTCTTCTGGGTGGCAGT-3’), for USP9x (F: 5’-AAGTGAAGCATGTC AGCGATT-3’, R: 5’-GCCACACATAGCTCCACCA-3’), for SOX2 (F:5’-GAGAACCCCAAGATGCACAAC-3’, R: 5’-CGCTTAGCCTCGTCGATGA-3’).

### Immunoprecipitation

HEK 293T cells were transfected with plasmids encoding pMSCV-SOX2-HA and pEZ-M12-Flag-USP9x. Forty-eight hours later, cells were lysed in 1 ml RIPA (50 mM Tris–HCl, pH 7.4, 150 mM NaCl, 1 mM EDTA, 1% Triton X-100, 2 mM NEM, cocktail (1:100)). Cell lysate was incubated with anti-Flag M2 beads or anti-HA M2 beads at 4 °C overnight, and beads were washed with RIPA (containing 2 mM NEM) 5 times. Samples were analyzed by immunoblotting.

### Plasmids

The shRNA or pGIPz plasmids, with the lentiviral packaging vectors PSPAX2 and pMD2G were introduced into HEK293T cells to produce lentivirus. After 48 h, the viral supernatant was collected and added to cells in six-well plates with medium containing 8 μg/ml polybrene (Sigma). After 2 days’ infection, stably transfected cells were selected with puromycin. The targeting sequences of shUSP9x were shown to be the following: shUSP9x-1^#^: 5’-CTTAAATCCTCATTGCAAA-3; 5^#^: 5’-GATGTATTCTC AATCGTAT-3’. For overexpressing SOX2, the target fragment of SOX2 was cloned from pMXs-hSOX2 with primers F: 5’-GCCTCGAGATGGATTACAAGGATGACGA CGATAAGATGTACAACATGATGGAGAC-3’, R: 5’-GCGAATTCTCACATGTGTGAGAGGGGCAG-3’. the PCR product was cut with XhoI and BamHI, these fragments were ligated into pMSCV-puro linearized with XhoI/BamHI.

### Immunofluorescence analysis

The cells on glass coverslips were fixed with 4% paraformaldehyde, treated with 0.3% Triton X-100, and blocked with 2% bovine serum albumin. Cells were then sequentially incubated with antibody USP9x (Santa Cruz Biotech, sc-365353) or SOX2 (Abcam, ab97959) overnight at 4 °C, followed by FITC labeled anti-mouse immunoglobulin G antibody (Invitrogen) and Cy5.5 labeled anti-rabbit immunoglobulin G antibody (Invitrogen). Stained cells were examined with immunoflurescence microscopy (Nikon).

### Xenograft studies

Five-week-old male nude mice (BABL/c) were purchased from Shanghai Slack Laboratory Animal Co., LTD, and maintained in a standard environment. Per mice was injected subcutaneously in the left or right flank with 2×10^6^ 143B cells suspended in 100 μl PBS. Tumors were allowed to reach about 100 mm^3^, after which mice were tumor-size matched and were treated with NGA (i.p. 6mg/kg) or vehicle (DMSO) every other day. Tumor size was monitored by calipers every other day using the following formula: volume= 1/2×length×(width)^2^.

## Results

### NGA inhibits proliferation and induces apoptosis in osteosarcoma cells

To explore the antitumor effects of NGA in osteosarcoma, osteosarcoma cell lines U2OS and 143B were treated with NGA for 24 h or 48 h, respectively. Cell proliferation was evaluated by CCK8 assay (Fig 1B). NGA inhibited the proliferation of tumor cells with IC_50_ at 1∼2 μM. To investigate whether NGA induce apoptosis of osteosarcoma cells, U2OS and 143B were exposed to different doses of NGA for various times. NGA treatment resulted in the cleavage of caspase-3 and PARP1 in U2OS and 143B in a dose and time-dependent manner (Fig 1C). Consistent with these observations, NGA increased the percentage of apoptotic cells, as evaluated by Annexin V/PI staining (Fig 1D). Moreover, NGA effectively inhibited the formation of colony in various cells (Fig 1E), as examined with crystal violet staining. These data indicated that NGA could inhibit proliferation and induce apoptosis of osteosarcoma cells.

**Figure 1.**
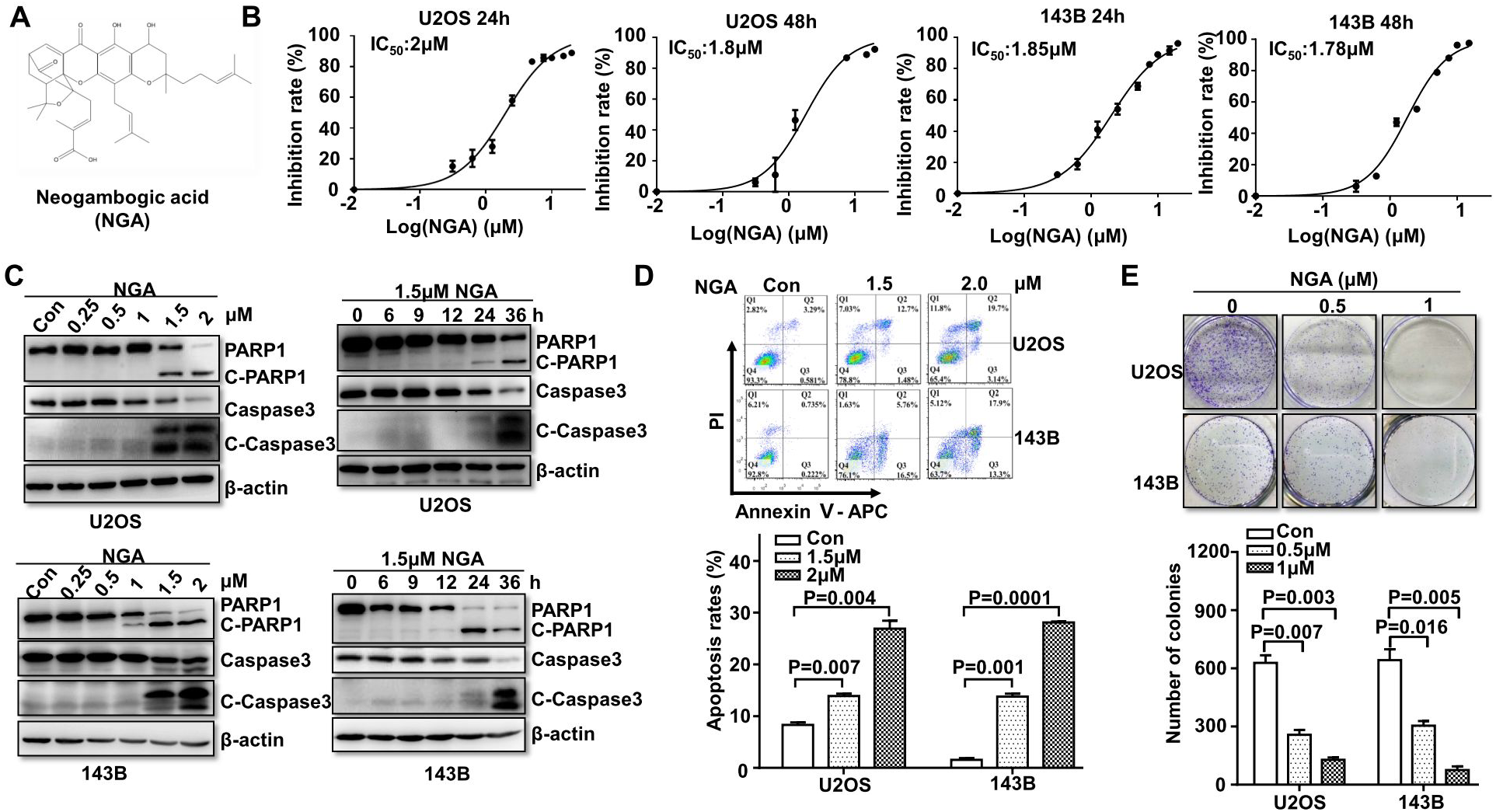
NGA inhibits the proliferation, induces apoptosis and inhibits colony formation in osteosarcoma cells. A Chemical structure of NGA. B Osteosarcoma cell lines were treated with different doses NGA for 24 h or 48 h. Cell proliferation was examined by CCK8 assay and the IC_50_ of NGA were calculated. C The osteosarcoma cells U2OS and 143B were treated with different doses of NGA for 24 h or 1.5 μM NGA for different times, and the indicated proteins were examined by western blot. D The cells were exposed to different concentration of NGA for 24h, and then Annexin/PI double staining were used to evaluate the apoptosis rates. E Osteosarcoma cells were seeded into 6-well plates at a density of 2000 cells/well and cultivated for 10 days with 0.5 μM or 1 μM NGA, and then colony formation was detected by crystal violet staining.

### Ubiquitination-mediated degradation of SOX2 contributes to the anti-osteosarcoma effect of NGA

Since SOX2 plays a key role in the proliferation of osteosarcoma, we hypothesized that NGA may affect the expression of SOX2. Interestingly, NGA could reduce the protein level of SOX2 in a dose- and time-dependent manner (Fig 2A). The decrease of was proteasome dependent, as the proteasome inhibitor MG132 could remarkably reverse NGA-induced reduction of SOX2 (Fig 2B) and NGA did not decrease the mRNA level of SOX2 (Fig 2C). To further confirm this, we examined the influence of NGA on the ubiquitination status of SOX2. We found that NGA treatment could obviously increase ubiquitination of SOX2 in cells, especially K48-linked ubiquitin, which closely related to proteasome-mediated protein degradation (Fig 2D). To test whether SOX2 contributed to the anticancer effect of NGA, SOX2 was overexpressed in U2OS and 143B cells (Fig 2D). The results showed that overexpression of SOX2 significantly weaken NGA-induced cell death in osteosarcoma cells (Fig 2E). These data indicated that degradation of SOX2 plays an important role in the anti-osteosarcoma effect of NGA.

**Figure 2.**
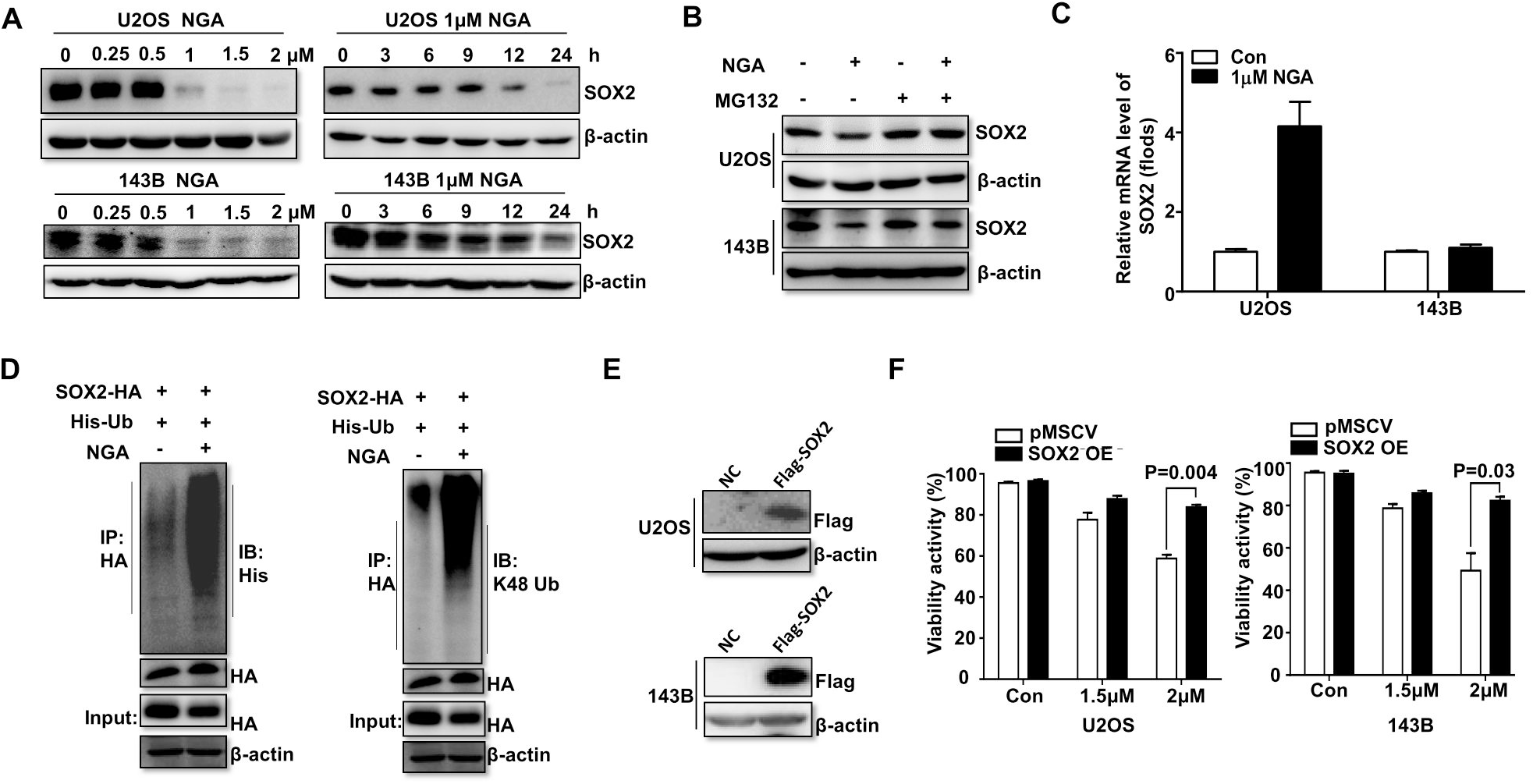
Ubiquitin proteasome pathway-mediated degradation of SOX2 contributes to the anti-cancer effect of NGA. A U2OS and 143B cells were treated with NGA with various concentrations for 24 h or 1 μM NGA for various times, and protein level of SOX2 was examined by western blot. B U2OS and 143B cells were treated with 1μM NGA for 12 h in the absence or presence of MG132 (5 μM, added at 6 h) and the indicated proteins were examined by western blot. C U2OS and 143B cells were treated with 1 μM NGA for 12 h, the mRNA level of SOX2 was examined by qPCR. D The plasmids SOX2-HA and His-Ub were co-transfected into HEK 293T cells, and then these cells were treated with NGA. Immunoprecipitation was performed with anti-HA antibody and the indicated proteins were examined by western blot. E,F SOX2 stably transfected U2OS and 143B cells (E) were treated with NGA and cell viability was measured (F).

### NGA inhibits osteosarcoma growth and reduces SOX2 expression *in vivo*

In order to further demonstrate the anti-cancer of NGA *in vivo*, we used a xenograft mouse model. Compared to the control group, intraperitoneal administration of NGA (6 mg/kg) significantly decreased the size (Fig 3A and B) and weight (Fig 3C) of tumors formed by 143B cells. This could be due to inhibition of cell proliferation and an increase in cell apoptosis, as revealed by a decrease in Ki-67 staining and an increase in TUNEL positive cells. Consistent with the effect of NGA on SOX2 *in vitro*, NGA treatment also reduced the expression of SOX2 *in vivo* (Fig 3D). In addition, the mice were tolerant to NGA at the concentration used despite a slight fall of body weight was also observed (Fig 3E). These results indicated that NGA suppresses osteosarcoma growth and reduced SOX2 expression *in vivo*.

**Figure 3.**
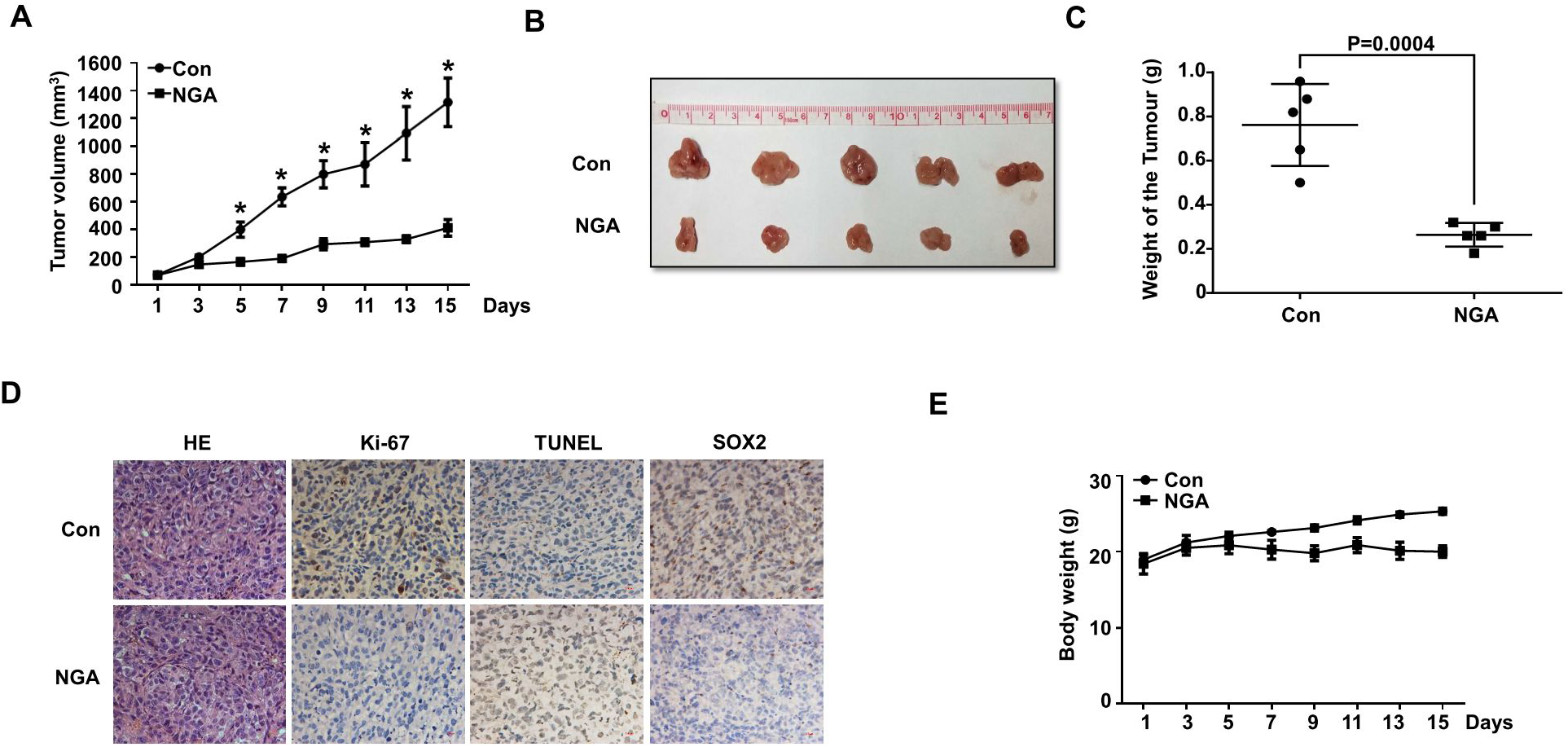
NGA inhibits osteosarcoma growth and reduce SOX2 expression *in vivo*. A 143B xenografts in nude mice were treated with vehicle or NGA (every other day by intraperitoneal injection) for 15 days, tumor volumes were recorded. Values are expressed as the mean ± SD. *\*P<0.05* vs. control group. B,C Image of xenograft tumors treated with NGA or vehicle on day 15 (B) and tumor weights (C) were shown. D The expression patterns of Ki-67, TUNEL, and SOX2 were tested by immunohistochemistry analysis in the xenograft tumors on day 15 in each group. Original magnification, ×400. E Effect of NGA on mouse body weight.

### USP9x maintains SOX2 stability

The increase of SOX2 ubiquitination maybe due to activation of E3 ubiquitin ligase or inactivation of DUBs for SOX2. Generally, small molecules such as NGA are more likely to function as an inhibitor than an activator. We hypothesized that NGA may inhibit the DUB for SOX2. However, the DUB for SOX2 is currently unknown. Therefore, we first performed a screen to identify the DUB for SOX2. Thirty DUBs were overexpressed in HEK 293 cells and their effect on the protein level of endogenous SOX2 was analyzed by western blot. We found that USP9x remarkably upregulated SOX2 protein level (Fig 4A). It seemed that some other DUBs such as USP7, USP2, USP14 had effect on SOX2. However, the commercial available inhibitors for these DUBs had no obvious influence on the SOX2 expression, including b-AP15 (the inhibitor of USP14 and UCHL5), P5091 (the inhibitor of USP7 and USP47) and ML364 (the inhibitor of USP2), except WP1130 (the inhibitor of USP9x) (Fig S1). To confirm USP9x regulate the stability of SOX2, USP9x was overexpressed in cells. Overexpression of USP9x resulted in SOX2 elevation in a dose-dependent manner (Fig 4B), and significantly prolonged the half-life of SOX2 (Fig 4C). In contrast, depletion of USP9x by shRNA (Fig 4D) or inhibiting USP9x activity by WP1130 (Fig 4F) could markedly decrease SOX2 protein (Fig 4E and F) but not its mRNA level (Fig S2A and B). Moreover, the decrease of SOX2 was reversed by addition of proteasome inhibitor MG132 (Fig 4E and F). These results demonstrated that USP9x specifically stabilizes SOX2 in cells.

**Figure 4.**
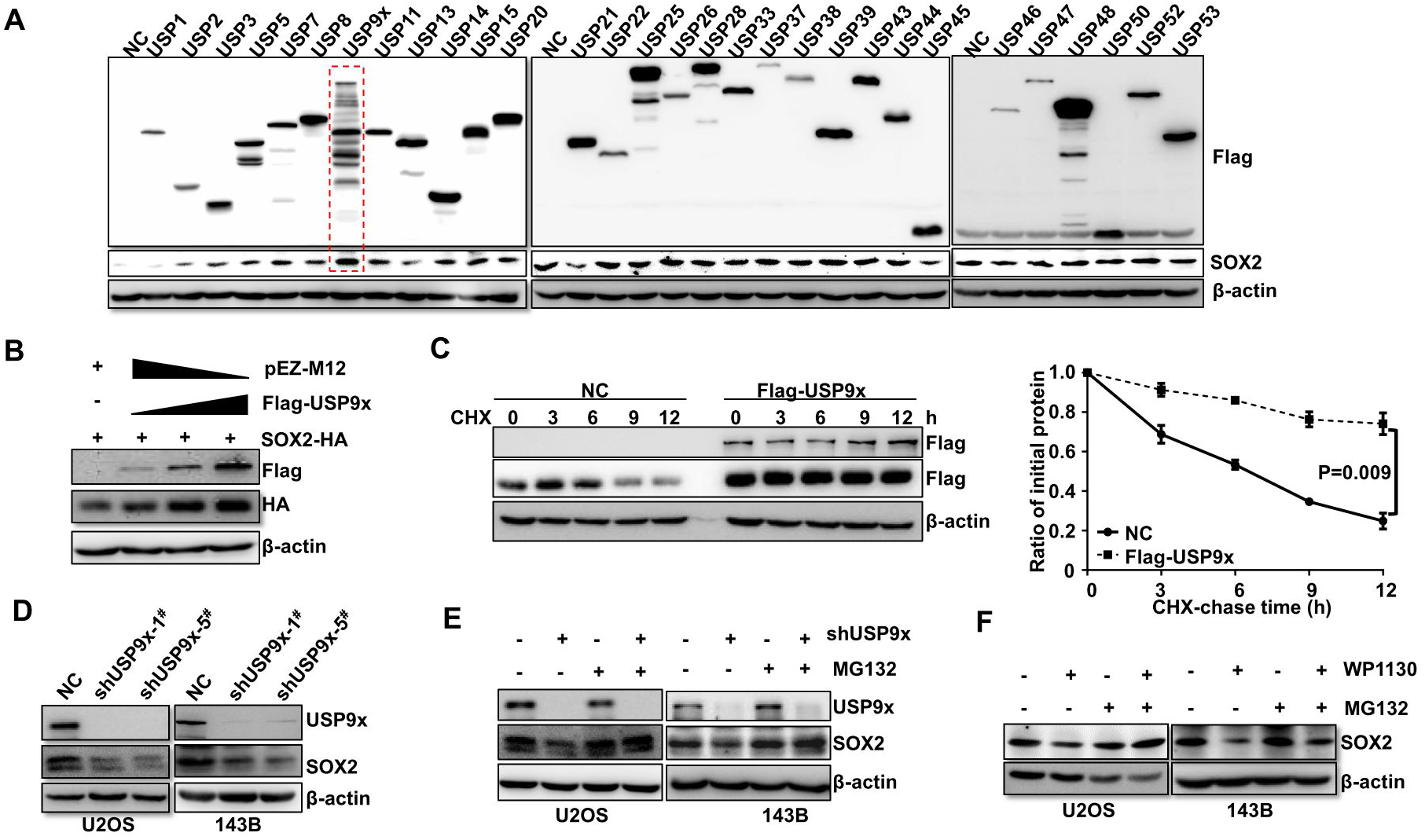
USP9x stabilize SOX2 protein. A The indicated DUBs were transfected into 293T cells, respectively. Forty-eight hours later, cells were harvested and the indicated proteins were examined by western blot. B Increasing amounts of USP9x plasmids were co-transfected with SOX2 plasmids into 293T cells and the indicated proteins were examined by western blot. C 293T cells transfected with USP9x and SOX2 plasmids were treated with cycloheximide (10 µg/ml), and collected at the indicated times for western blot (left). Quantification of SOX2 levels relative to β-actin was shown (right). Results were shown as mean ± s.d. D USP9x was depleted by shRNAs in the cell lines and the indicated proteins were examined by western blot. E USP9x silenced or the control cells were treated with or without MG132 and the indicated proteins were examined by western blot. F Osteosarcoma cells were treated with USP9x inhibitor WP1130 in the presence or absence of MG132 and the indicated proteins were examined by western blot.

### USP9x interacts with and deubiquitinates SOX2

If SOX2 is the substrate of USP9x, USP9x should interact with and regulate its ubiquitination. To confirm this hypothesis, we examined the interactions between exogenous SOX2 and endogenous USP9x, or between exogenous SOX2 and exogenous USP9x. USP9x and SOX2 were readily co-immunoprecipitated with each other (Fig 5A and B). To further demonstrate their interaction, the co-localization of USP9x and SOX2 in osteosarcoma cells was detected by immunofluorescence. As shown in Fig 5C, co-localization of USP9x and SOX2 was observed in osteosarcoma cells. Next, we examined whether USP9x could regulate the ubiquitination of SOX2. As expected, knockdown of USP9x increased SOX2 ubiquitination in cells (Fig 5D left) while ectopic expression of USP9x decreased K48-but not K63-linked ubiquitination of SOX2 (Fig 5D). These data suggested USP9x indeed regulates the ubiquitination of SOX2 in cells.

**Figure 5.**
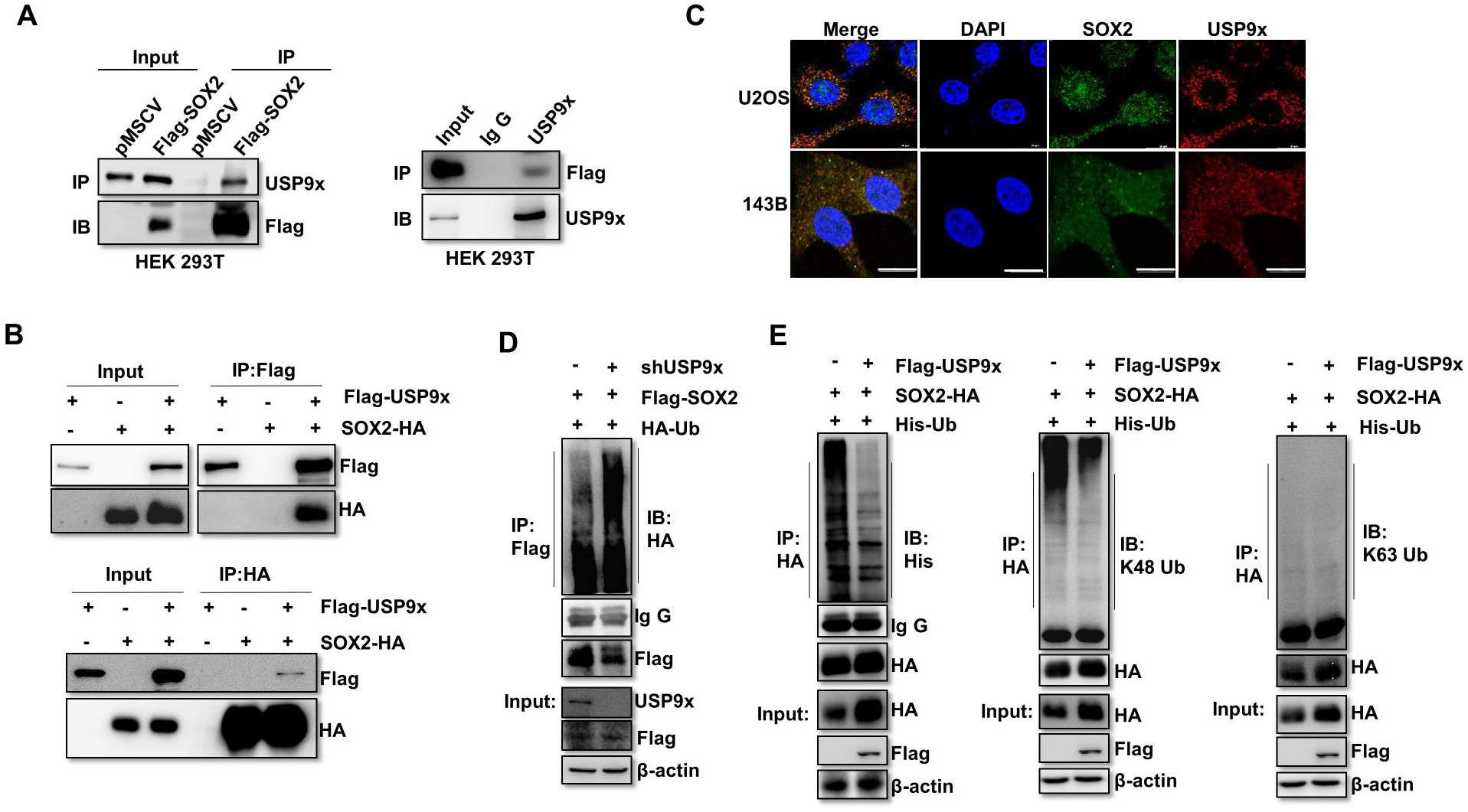
USP9x interacts with and de-polyubiquitylates SOX2. A HEK 293T cells were transfected with Flag-SOX2 plasmids, and immunoprecipitation (IP) was performed using Flag-M2 beads or USP9x antibody. The indicated proteins were examined by western blot. B Cell lysates from HEK 293T cells transfected with Flag–USP9x and SOX2-HA plasmids were subject to immunoprecipitation (IP) with anti-Flag or anti-HA antibodies. The lysates and immunoprecipitates were analyzed by western blot with the indicated antibodies. C Immunofluorescence staining of U2OS and 143B showing co-localization of USP9x (green) and SOX2 (red). DAPI (blue) was used for nuclear staining. D HEK 293T cells were transfected with HA-Ub, Flag-SOX2 and shUSP9x, immunoprecipitation (IP) was performed using Flag-M2 beads. The indicated proteins were examined by western blot. E HEK 293T cells were transfected with His-Ub, SOX2-HA and Flag-USP9x, IP was performed using HA antibody. The indicated proteins were examined by western blot.

### NGA interacts with USP9x in cells

Having demonstrated that USP9x is the DUBs for SOX2, we assumed that NGA may directly interact with USP9x in cells. To this end, the CETSA assay was performed. This method has been used to evaluate the possible interaction of a small molecule with its target protein in its native environment. As shown in figure 6A, NGA increased the thermal stability of USP9x with different temperature. Moreover, the effect of NGA on the thermal stability of USP9x was dose dependent (Fig 6B). This data indicated that NGA directly interacts with USP9x in cells.

**Figure 6.**
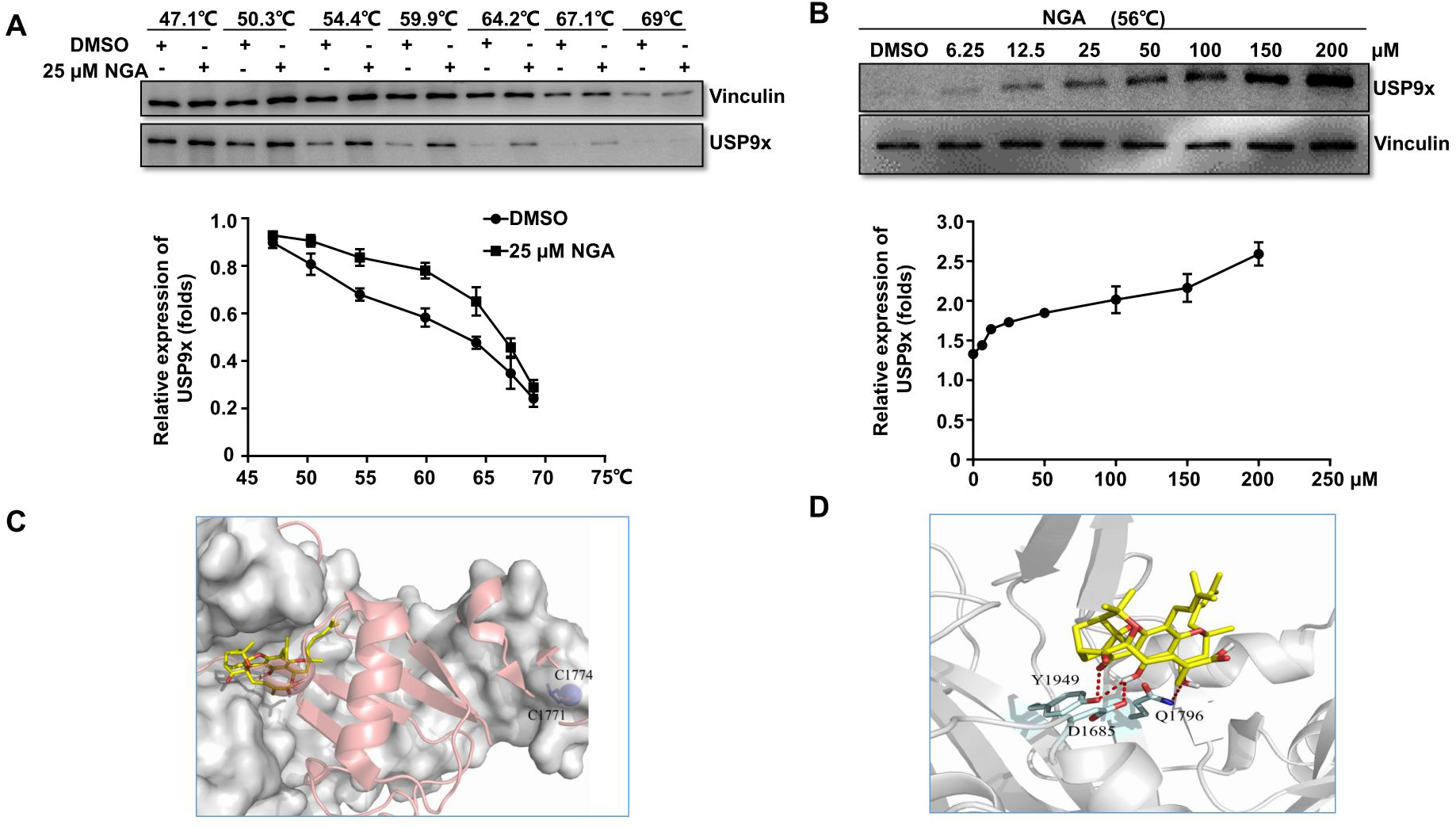
NGA interacts with USP9x in cells. A,B CETSA was performed on U2OS cells as described in materials and methods. The thermal stabilization effect of NGA on USP9x and vinculin at different temperatures (A) and different doses (B) were evaluated by western blot. The intensity of the bands was quantified by Image J software. All experiments were repeated three times with the same results. C Superposition of the USP9x and USP7/Ub-aldehyde (PDB ID code 1NBF) structures. The molecular surface of USP9x is shown in gray. USP7/Ub-aldehyde was shown in pink cartoon. Predicted conformation of NGA in the binding pocket of USP9x was shown in yellow sticks. Zinc molecule was shown in blue sphere. D Residues provided for interactions with NGA in USP9x are shown in green sticks. The dashed lines in red represent hydrogen bonds.

Molecular docking was carried out to explore the binding mechanisms between NGA and USP9x. The result demonstrated that NGA binds to the same subpocket that can be occupied by the C-terminal tail of Ub (Figure 6C), which was different from compound WP1130. WP1130 was reported to block USP9x activity by forming reversible covalent adducts with Cysteines such as Cys1771 and Cys1774 (Figure 6C). Residue Tyr 1949, Asp1685 and Gln1796 forms polar interactions with NGA (Figure 6D). This result suggested that the inhibition mechanism of this compound might be explained by the blockage of Ub.

### Knockdown of USP9x induces proliferation decrease in osteosarcoma cells

We have demonstrated NGA can regulate the stability of SOX2 by directly inhibiting USP9x. We next tried to determine the role of USP9x in osteosarcoma. USP9x was silenced by shRNA in U2OS and 143B cells and the knockdown efficiency was accessed by western blot (Fig 7A). Knockdown of USP9x significantly suppressed the proliferation of osteosarcoma cells (Fig 7B and C). Flow cytometry analysis showed that knockdown of USP9x partially block the cells at G2/M phase (Fig 7D). Consistent with these results, knockdown of USP9x also inhibited the colony formation of osteosarcoma cells (Fig 7E). Importantly, reintroduction of SOX2 into USP9x-depleted cells could reverse USP9x depletion-induced inhibition of colony formation (Fig 7F). These data suggested that USP9x plays an oncogenic role in osteosarcoma cells.

**Figure 7.**
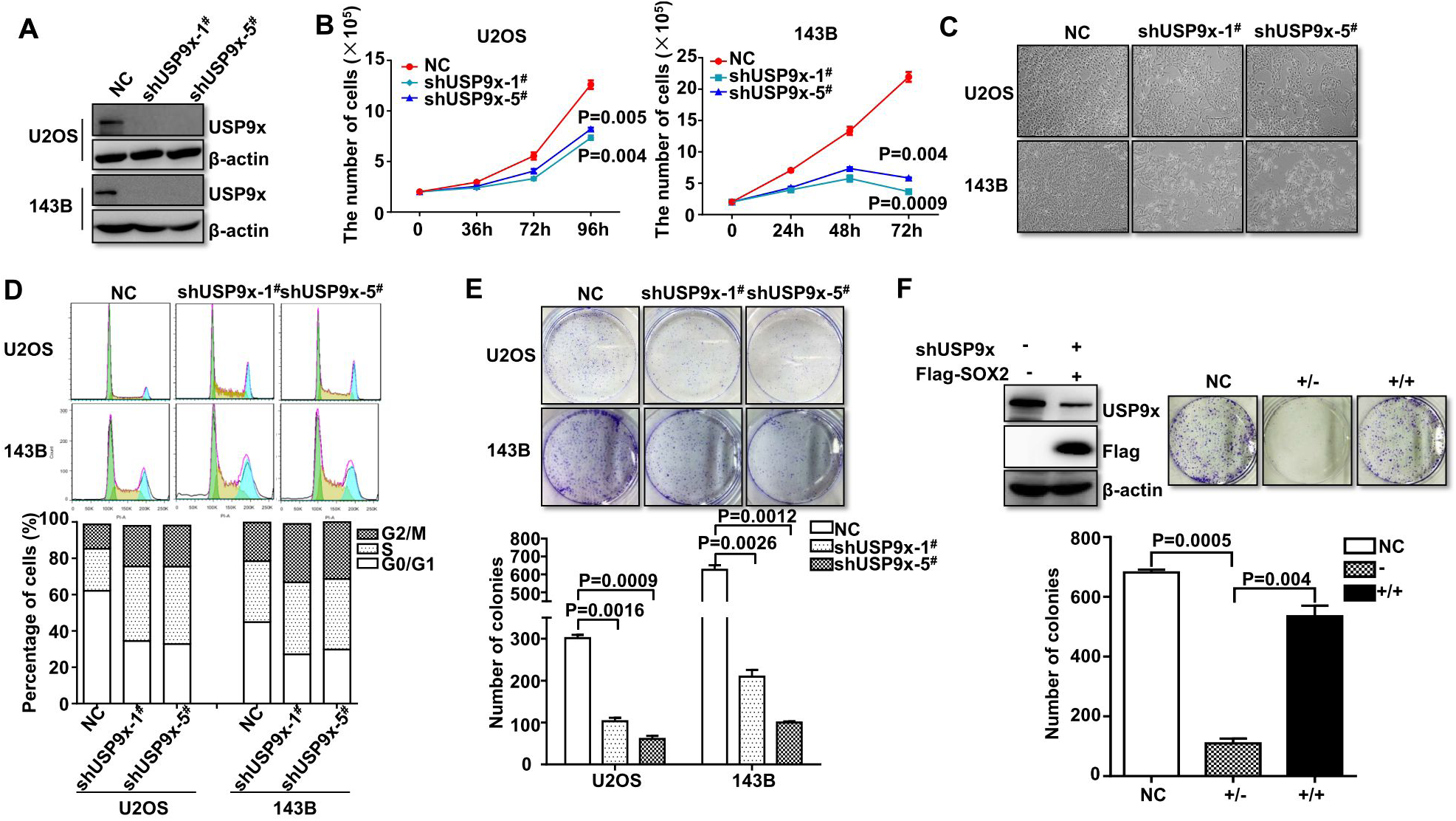
Knockdown of USP9x inhibits proliferation of osteosarcoma cells. A The USP9x specific shRNA or the non-specific control shRNA were transfected into U2OS and 143B cells, respectively. The indicated proteins were examined by western blot. B,C,D,E Cell proliferation of USP9x silenced and the control cells were monitored by trypan blue exclusion assay (B). Cell morphology (96 h or 72 h) were captured under the microscope (C). Cell cycle distribution was monitored by FACS (D). Clone formation capacity was detected by crystal violet staining assay (E). F 143B cells were transfected with shUSP9x with or without co-transfection of Flag-SOX2. Clone formation capacity was detected by crystal violet staining assay.

### Knockdown of USP9x suppresses proliferation of osteosarcoma cells *in vivo*

In order to investigate the anti-tumor effect of USP9x knockdown *in vivo*, a xenograft nude mouse model was established with subcutaneous inoculation of USP9x silenced 143B cells. The efficiency of USP9x knockdown in 143B cells was assessed by western blot analysis (Fig 8A). Compared to the control group, knockdown of USP9x markedly reduced the tumor size (Fig 8B) and weight (Fig 8C and D). Compared to control group, the immunohistochemistry assay showed that USP9x-silenced group has higher percentages of apoptotic cells, lower percentages of proliferative cells and lower level of SOX2 protein (Fig 8E). These results suggested that knockdown of USP9x inhibits proliferation of osteosarcoma cells with reduction of SOX2 *in vivo*.

**Figure 8.**
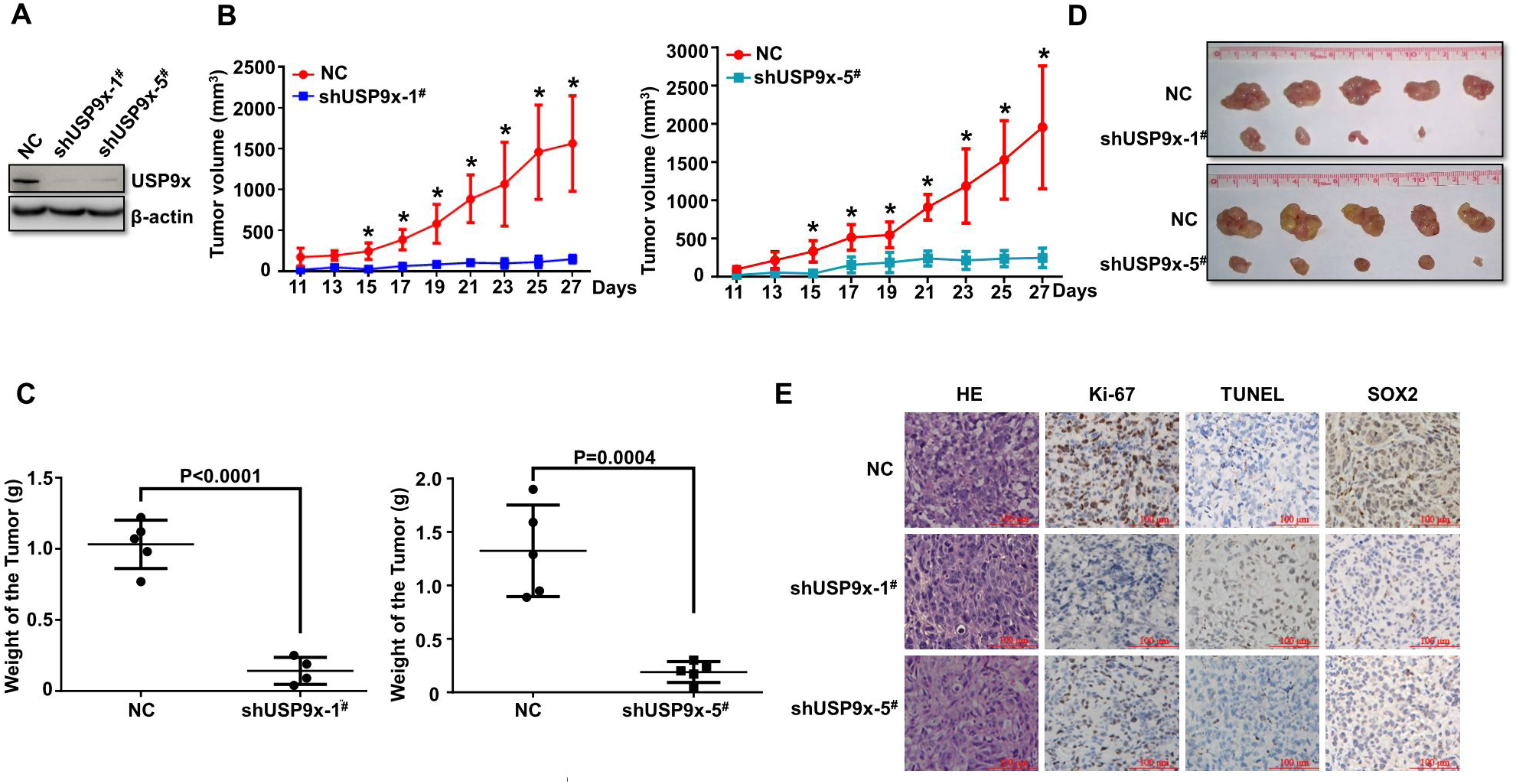
Knockdown of USP9x suppresses tumorigenicity of osteosarcoma cells. A The efficiency of knockdown of USP9x was examined by western blot. B,C,D,E 143B cells were inoculated subcutaneously into nude mice. Tumor volume was monitored every other day when it reached 100 mm^3^ (B), the symbols * indicated *P<0.05*. Tumor weights were measured on day 27 (C), and the representative images of xenograft tumor were shown (D). (E) Expression patterns of Ki-67, SOX2 and TUNEL positive cells examined by immunohistochemistry in the xenograft tumors in each group. Original magnification, × 400.

## Discussion

Osteosarcoma is the most common primary malignant bone cancer (Gianferante et al, 2017). Despite the 5-year survival rate increased over the past 40 years, no substantial improvement has been obtained since the 1980s (Kansara et al, 2014; Tiram et al, 2016). It is required to find novel target to combat this disease. In this study, we demonstrate that inducing degradation of SOX2 by inhibiting USP9x is a promising strategy to combat osteosarcoma. And NGA is a novel USP9x inhibitor for the treatment of osteosarcoma.

Many transcription factors have been involved in the pathogenesis of cancer and were considered as drug targets. However, different from that of protein kinases, it is difficult to develop direct transcription factor “inhibitors”. An alternative strategy is to inducing the degradation of these transcription factors. Both ubiquitin-proteasome and autophagy pathway have been involved in the regulation of SOX2 turnover. Theoretically, activating the E3 ligase or inhibiting the DUBs could lead to protein degradation. Ub-conjugating enzyme E2S (UBE2S), has been reported as the E3 ubiquitin ligase for the ubiquitination SOX2 (Fang et al, 2014; Wang et al, 2016; Wang et al, 2019; Zhang et al, 2019). However, develop an agonist for E3 ligase is more difficult than develop an antagonist for DUBs. In fact, several successful DUBs has been used to induce the degradation of transcription factors (D’Arcy et al, 2011). For example, P22077, a selective and potent inhibitor of ubiquitin-specific protease 7 (USP7), has been used to induce the degradation of N-MYC and NOTCH1 in neuroblastoma and acute lymphoblastic leukemia (Chauhan et al, 2012; Tavana et al, 2016). Therefore, we attempt to induce the degradation of SOX2 by targeting its DUBs.

However, despite that USP22 has been shown to repress the transcription of SOX2, the deubiquitinating enzymes for controlling stability of SOX2 is currently not known (Sussman et al, 2013). In this study, we identified USP9x as a bona fide DUBs for SOX2. Indeed, USP9x can interact with and increase the stability of SOX2 through regulating the K48-linked ubiquitination of SOX2. Moreover, we found that knockdown of USP9x or using USP9x inhibitor can inhibit cell proliferation, which could be rescued by overexpression of SOX2. Therefore, we proposed that inducing the degradation of SOX2 by targeting its deubiquitinating enzyme is a promising strategy to combat osteosarcoma. In searching for the DUBs for SOX2, it seemed that several other DUBs such as USP2, USP7, and USP14 also have some effect on the protein level of SOX2. However, because the inhibitors for USP2, USP7 or USP14 cannot decrease the protein level of SOX2, it is unlikely that these DUBs are the DUBs responsible for SOX2 stability. As this a small scale of screening (compared to the total 100 DUBs, only 30 of them were used in our test), we don’t rule out other DUBs may be also involved in the regulation of SOX2 stability.

The function of USP9x is cell-type specific. USP9x has been viewed as a tumor suppressor gene in colorectal and pancreatic ductal adenocarcinoma, while others reported that USP9x is an oncogene in breast cancer, glioblastoma, and leukemia (Akiyama et al, 2019; Chen et al, 2019; Khan et al, 2018a; Pal et al, 2018; Potu et al, 2017b; Zhu et al, 2018). Our results suggest that USP9x may function as an oncoprotein through regulating SOX2 in osteosarcoma. As expected, knockdown of USP9x or using USP9x inhibitors inhibits the proliferation of osteosarcoma cell lines *in vitro* and *in vivo*. Due to the short of osteosarcoma patient data for analysis, the relationship between the expression of USP9x and the prognosis of osteosarcoma is not clear. Future analysis of large amount of patient data of osteosarcoma is needed.

Currently, the known USP9x inhibitor is WP1130 (Liu et al, 2015b). WP1130 has shown great activity in a variety of cancers including chronic myelogenous leukemia, FLT3-ITD positive leukemia, myeloma, glioblastoma, liver cancer and lung cancer (Bartholomeusz et al, 2007; de Las Pozas et al, 2018; Kapuria et al, 2010; Kim et al, 2019; Liu et al, 2015b). Since NGA could also inhibit USP9x, it is likely NGA also has some activity against these cancers. To date, NGA do show anti-cancer activity against melanoma, myeloma and leukemia. Interestingly, the mode of action of NGA on USP9x may be different from that of WP1130. WP1130 treatment leads to the reduction of USP9x protein, while NGA does not lead to the degradation of USP9x (Fig S3), suggesting a novel mechanism of inhibition (Peterson et al, 2015). Our molecular docking studies support this notion, WP1130 and NGA bind to different sub-pockets of USP9x. In addition, both WP1130 and NGA are not very specific inhibitors for USP9x. Further investigation of the interaction between NGA and USP9x may provide information for optimizing more specific USP9x inhibitors.

In summary, we demonstrated that USP9x can deubiquitinate and stabilize SOX2 in osteosarcoma cells and USP9x is a novel therapeutic target for the treatment of osteosarcoma. NGA is a novel promising USP9x inhibitor for the treatment of osteosarcoma. Since SOX2 is involved in the regulation of embryonic stem cell and other cancer stem cells, the finding of USP9x/SOX2 axis may have broader significance than osteosarcoma.

## Acknowledgements

This work was supported in part by grants from the National Key Research and Development Program of China (No.2017YFA0505200), National Basic Research Program of China (973 Program) (NO. 2015CB910403), National Natural Science Foundation of China (81570118, 81670139, 81870156), Natural Science Foundation of Shanghai (19ZR1428700), Natural Science Foundation of Inner Mongolia Autonomous Region of China (2018MS08134) and State Key Laboratory of Bioorganic and Natural Product Chemistry.

## Author Contributions

YW, WG and XC designed the research; XC, XZ, HC, WY, HL, HX, WW, QZ, JK performed the research; XC, XZ, HC, WY, HL, HX, WW, QZ, JK analyzed data; Y W, WG, XC and XZ wrote the paper; YW,WG and TY supervised the study. All authors read and approved the manuscript.

